# Transition to siblinghood causes substantial and long-lasting physiological stress reactions in wild bonobos

**DOI:** 10.1101/2022.02.14.480345

**Authors:** Verena Behringer, Andreas Berghänel, Sean M. Lee, Barbara Fruth, Gottfried Hohmann

**Affiliations:** Endocrinology Laboratory, German Primate Center, Leibniz Institute for Primate Research, Göttingen, Germany; Max Planck Institute for Evolutionary Anthropology, Leipzig, Germany; Domestication Lab, Konrad Lorenz Institute of Ethology, Department of Interdisciplinary Life Sciences, University of Veterinary Medicine Vienna, Vienna, Austria; Primate Behavioral Ecology Lab, Center for the Advanced Study of Human Paleobiology, Department of Anthropology, George Washington University, Washington, USA; Max Planck Institute of Animal Behavior, Konstanz, Germany; Centre for Research and Conservation, Royal Zoological Society of Antwerp, Antwerp, Belgium

**Author notes:** These authors contributed equally to this work (Verena Behringer, Andreas Berghänel).

**Keywords:** Sibling birth, sibling rivalry, weaning, immature *Pan paniscus*, life history event, early life adversity, non-human primate, cortisol, neopterin, triiodothyronine, non-invasive

## Abstract

In mammals with a slow ontogeny, the birth of a sibling marks a major developmental transition. Behavioral studies suggest that this event is stressful for the older offspring, but physiological evidence for this is lacking, and it remains unknown whether the birth of a sibling is stressful beyond mere weaning stress. Studying transition to siblinghood in wild bonobos, we investigated physiological changes in urinary cortisol (stress response), neopterin (cell-mediated immunity), and total triiodothyronine (metabolic rate), and related them to behavioral changes in mother-infant relationship and feeding (suckling, riding, proximity, body contact, independent foraging). With sibling’s birth, cortisol levels increased fivefold in the older offspring and remained elevated for seven months, independent of age. This was associated with diminished immunity but not with behavioral or metabolic changes. Our results indicate that transition to siblinghood is stressful beyond nutritional and social weaning and suggest that this effect is evolutionary old.

## Introduction

Most children are exposed to the birth of a younger sibling while still being dependent on parental support. For the older offspring, this transition to siblinghood (TTS) marks the onset of considerable changes, including the sudden emergence of a competitor (sibling rivalry, (Dettwyler, 2017; Myers and Bjorklund, 2018)) and a decline in maternal investment (Kramer, 2011). Therefore, many researchers consider TTS to be a stressful life event or even disruptive crisis for the older sibling even under favorable conditions, a perspective that seems to be supported by behaviors such as aggression, clinginess, and depressive syndromes (reviewed in Volling, 2012; Volling et al., 2017). However, whether behavioral changes during TTS are the direct result of stressors associated with sibling birth rather than mere behavioral adjustments remains unclear (Volling, 2012; Volling et al., 2017), and TTS-related changes in physiological stress levels have to our knowledge never been measured in humans or other animals. Therefore, despite being a topic of intensive research in pediatrics, psychology, and social and health sciences, the assumption that TTS is a physiologically stressful life event or even crisis for the older sibling remains untested. In our study, we investigate for the first time TTS-related changes in physiological stress response (cortisol levels) in a wild ape (bonobos, *Pan paniscus*), as a referential model with a human-like TTS, and compare it with other behavioral and physiological data to test whether such changes are the result of sibling birth and/or simply the result of social and energetic effects associated with weaning.

Many human studies illustrate the sudden changes in older children upon sibling birth. The birth of another child induces mother-offspring conflict over resource allocation that can have lasting consequences for the older offspring (Trivers, 1974). Mothers allocate social attention and resources to the new born, and in humans, the quality and intensity of maternal interactions with the older sibling can decrease dramatically, inducing sibling rivalry (Baydar et al., 1997; Stewart et al., 1987). The psychological symptoms associated with TTS include e.g., anxiety, attention-seeking, social isolation, and *temper tantrums* (Baydar et al., 1997; Legg et al., 1974; e.g., Volling et al., 2017). However, recent studies revealed that such effects of TTS may instead relate to changes in the caregiving environment that sometimes accompany sibling birth, such as extended separation from the mother, rather than the birth of the sibling *per se* (Oh et al., 2017; Volling, 2012; Volling et al., 2017). The assumption that TTS in humans is stressful is primarily based on behavioral data from interviews, questionnaires or observations (e.g., Baydar et al., 1997; Legg et al., 1974). Quantitative physiological assessments of TTS-related stress responses in older siblings have never been conducted.

TTS-related stress responses could result from sibling rivalry, nutritional and social weaning from the mother, and /or other changes in the socioecological environment of the older offspring. In contrast to species such as rodents and birds in which sibling rivalry among same-aged litter mates can be severe or even lethal (Forbes, 2010), humans and non-human primates tend to give birth to singletons. Therefore, siblings differ in age and their dependence on mothers for food, transport, and social support, and thus, weaning status. In primates, weaning is a slow process (Langer, 2008), and particularly social dependence on the mother often lasts until the offspring reaches reproductive maturation (Lonsdorf et al., 2018; van Noordwijk et al., 2018). Accordingly, females of many species give birth to another infant before the older offspring reaches full independence, resulting in an overlap of dependent siblings of different age (Achenbach and Snowdon, 1998). In nonhuman primates, sibling birth affects the interactions between the older offspring and the mother (Schino and Troisi, 2001), which may affect the fitness of the older offspring throughout life. For example, shorter inter-birth intervals were related to reduced somatic growth in the older offspring in wild chimpanzees (Emery Thompson et al., 2016), and related to reduced fertility, weakened social connectedness and diminished survival of the older offspring in wild baboons (Alberts, 2019; Tung et al., 2016; Zipple et al., 2019).

TTS could overlap with and/or accelerate nutritional and social weaning, which on its own is known to be stressful in primates and other mammals (e.g., Hau and Schapiro, 2007; Mandalaywala et al., 2014). As a result, it is difficult to differentiate between the effects of sibling birth and weaning (Volling, 2012; Weary et al., 2008). However, humans and non-human apes have highly variable inter-birth intervals, such that sibling birth can occur at very different weaning stages, with some older offspring still nutritionally dependent on their mothers and others fully weaned for years at the time of sibling birth.

In our study, we leveraged the large variation in inter-birth intervals in wild bonobos to differentiate between the effects of TTS and age-related weaning and maturation. Apes offer a particularly suitable referential model to explore changes during TTS. They share a number of relevant traits with humans, like altricial state at the time of birth, extended social dependency from the mother, slow growth, late reproductive maturation, and long-life expectancy (Charnov and Berrigan, 1993; Jones, 2011). Additionally, maternal support is intense and persists for a long time (Stanton et al., 2020; van Noordwijk et al., 2018), and it is common that females give birth to another infant before the older offspring has reached social or even nutritional independence, resulting in an extended period of co-residence of two dependent offspring of different ages (Achenbach and Snowdon, 1998).

Humans and bonobos show similar patterns of physiological maturation in terms of the emergence and timing of a juvenile pause and adrenarche (Behringer et al., 2014, 2012). In bonobos, the onset of sexual maturation starts around seven years of age in males and four years in females (Behringer et al., 2014), who reach menarche at about 8.2 years (Thompson-Handler, 1990). In the wild, females emigrate from their natal groups at seven to nine years and give birth for the first time at an estimated age of 13-15 years (Furuichi, 1989; Hashimoto, 1997; Kuroda, 1989). Bonobos depend heavily on their mothers during the first two years of life, being usually in close contact and mainly carried (De Lathouwers, 2004; Kuroda, 1989; Lee et al., 2020). After the age of five years, spatial distance to the mother increases (Kuroda, 1989; Toda et al., 2021). Nutritional weaning is largely completed between four to five years (Kuroda, 1989; Oelze et al., 2020). Behavioral observations suggest that weaning is less stressful in bonobos than in chimpanzees (de Lathouwers and Van Elsacker, 2006), which is further supported by a lack of change in urinary cortisol around weaning age in immature bonobos (Tkaczynski et al., 2020). General changes in cortisol levels hint on physiological responses to the birth of a sibling instead (Tkaczynski et al., 2020).

Inter-birth intervals ranged from 2.3 to 8.6 years in our study population (average ± SD: 5.4 ± 1.5years) (Tkaczynski et al., 2020), and the state of the older siblings at the time when mothers gave birth to another infant ranged from highly dependent to rather independent. This wide variation in inter-birth intervals enables us to investigate for the first time the specific effects of TTS and offspring age on a) the physiological stress response, b) immunity, c) energetic stress, d) changes in the relationship between mothers and the older offspring and e) changes in foraging and travelling independency, while accounting for f) offspring sex and temporal environmental changes. This comprehensive approach in a species with a slow human-like ontogeny, living under natural conditions allows us to start to disentangle stress response patterns of TTS and weaning.

We measured urinary cortisol to assess stress responses before and during TTS. Changes in cortisol provide a physiological marker to quantify stress responses in humans and other mammals. Cortisol is produced in response to physical as well as psycho-social stressors (Kirschbaum and Hellhammer, 1994; McEwen, 2009). In children, salivary cortisol levels increase during traumatic family events (Flinn et al., 2012), indicating that cortisol measurements are a valuable tool to assess children’s stress responses to family interactions (Flinn et al., 2012) and thus, also TTS. Hence if TTS is a stressor, we expected a sudden increase in cortisol levels at sibling birth. Using aliquots of the same urine samples, we also measured neopterin as a marker of cell-mediated immunity and total triiodothyronine (total T3) as a proxy for metabolic changes. T3 is a thyroid hormone that increases metabolic rate, allowing us to disentangle the effect of energetic and social stressors which may both occur around the age of weaning (Maestripieri, 2018; Mandalaywala et al., 2014). If TTS affects the homeostasis of the older offspring, we expect a decline in neopterin levels with the birth of a sibling, and if sibling birth causes metabolic issues for the older offspring, we expect a decline in total T3 levels at sibling birth.

Physiological measures were complemented with behavioral scores on nursing and riding time, time in body contact and five-meter proximity to the mother as well as independent foraging. These parameters indicate pattern of social and nutritional weaning around TTS. We compared trajectories around sibling birth of these parameters to investigate whether TTS related changes in cortisol could be linked to similar changes in our other variables and, thus, to TTS-related weaning patterns.

All our response variables naturally also change with age. Therefore, age-related changes might a) directly mediate potential changes during TTS in case of strong temporal overlap, and/or b) moderate these effects as the importance of a TTS might decline with decreasing dependency on the mother. Therefore, for all our response variables, we first run a model with age as a control variable to control for potential mediation, followed by a model with age as an interaction term to further allow for such moderation effects (for more details see methods section). If TTS has effects beyond age-related weaning, then we expect sudden changes at sibling birth also after controlling for age-related changes.

## Results

Our main results are summarized in Table 1, and model structures can be derived from Table 2 and 3.

**Table 1:** Summary of the main findings in physiological and behavioral marker in the older offspring in relation to TTS.

|  | Cortisol | Neopterin | Total T3 | Suckling | Riding | 5m-proximity<br>with mother | Body contact<br>with mother | Independent<br>foraging |
| --- | --- | --- | --- | --- | --- | --- | --- | --- |
| **Sudden change at sibling**<br>**birth** | **Yes, increase** | **Yes, decrease** | *No* | *No* | **Yes, decrease** | *No* | **Yes, increase** | *No* |
| **Effect of TTS attenuates**<br>**with offspring age** | *No* | *No* | *No* | *No* | **Yes, effect**<br>**exists only up**<br>**to 5yrs old** | *No* | **Yes** | *No: all changes*<br>*finished before*<br>*sibling birth* |

**Table 2:** GAMM results for physiological changes (urinary cortisol, urinary neopterin and urinary total T3 levels) in the older offspring around sibling birth. Green: Classic interaction term derived from a separate model calculation (see methods section). ID: Individual. T3 = total T3. S-birth = sibling birth, “:” = interaction term. * before = before sibling birth, * after = after sibling birth, * early after = seven months following sibling birth, * late after = time following early after.

| Reference |  | log(Cortisol) |  |  |  | log(Neopterin) |  |  |  | log(T3) |  |  |  |
| --- | --- | --- | --- | --- | --- | --- | --- | --- | --- | --- | --- | --- | --- |
| Factor Variables: | Category | Est. | SE | t | p | Est. | SE | t | p | Est. | SE | t | p |
| (Intercept) |  | 0.99 | 0.05 | 18.80 |  | 2.41 | 0.04 | 57.54 |  | 0.89 | 0.06 | 15.97 |  |
| Males | Females | 0.10 | 0.06 | 1.73 | 0.083 | -0.03 | 0.04 | -0.70 | 0.49 | -0.11 | 0.06 | -1.77 | 0.078 |
| After S-birth* | Before* | - | - | - | - | -0.19 | 0.06 | -2.91 | **0.004** | 0.13 | 0.08 | 1.59 | 0.11 |
| Early after S-birth* | Before* | 0.67 | 0.09 | 7.4 | **< 0.001** | - | - | - | - | - | - | - | - |
| Early after S-birth* | Late after* | 0.60 | 0.09 | 6.47 | **< 0.001** | - | - | - | - | - | - | - | - |
| Late after S-birth* | Before* | 0.08 | 0.09 | 0.91 | 0.36 | - | - | - | - | - | - | - | - |
| Smooth term variables: |  | edf | Ref.df | F | p | edf | Ref.df | F | p | edf | Ref.df | F | p |
| Time-S-birth: males |  | 1.00 | 1.00 | 2.06 | 0.15 | 1.00 | 1.00 | 0.50 | 0.48 | 1.00 | 1.00 | 4.06 | **0.049** |
| Time-S-birth: females |  | 1.00 | 1.00 | 2.27 | 0.13 | 1.00 | 1.95 | 0.62 | 0.56 | 1.00 | 1.00 | 0.01 | 0.92 |
| Time-S-birth: males | Females | 1.00 | 1.00 | 0.03 | 0.86 | 1.00 | 1.00 | 0.30 | 0.59 | 1.00 | 1.00 | 3.10 | 0.078 |
| Age: males |  | 1.00 | 1.00 | 2.31 | 0.13 | 3.31 | 4.01 | 4.04 | **0.003** | - | - | - | - |
| Age: females |  | 1.00 | 1.00 | 0.91 | 0.34 | 1.00 | 1.00 | 0.03 | 0.88 | - | - | - | - |
| Age: males | Females | 1.00 | 1.00 | 0.19 | 0.66 | 2.23 | 2.73 | 1.87 | 0.25 | - | - | - | - |
| Daytime |  | 1.00 | 1.00 | 45.48 | **< 0.001** | 1.00 | 1.00 | 4.80 | 0.029 | 2.11 | 2.57 | 1.77 | 0.14 |
| Seasonal effect |  | 2.30 | 3.00 | 8.85 | **< 0.001** | 0.52 | 3.00 | 0.22 | 0.28 | 0.00 | 3.00 | 0.00 | 0.72 |
| Random effects: |  |  |  |  |  |  |  |  |  |  |  |  |  |
| Time-S-birth per ID (smooth) |  | 0.00 | 111.0 | 0 | 0.24 | 0.00 | 110.0 | 0 | 0.28 | 7 | 106.0 | 0 | 0.010 |
| Age per ID (smooth) |  | 3.42 | 104.0 | 0 | 0.15 | 0.00 | 108.0 | 0 | 0.33 | - | - | - | - |
| Mother ID (intercept) |  | 0.00 | 13.0 | 0 | 0.24 | 0.00 | 13.0 | 0 | 0.29 | 0 | 13.0 | 0 | 0.21 |
| Year (intercept) |  | 0.00 | 1.0 | 0 | 0.060 | 0.00 | 1.0 | 0 | 0.29 | 0 | 1.0 | 0 | 0.88 |
| R<sup>2</sup><sub>adj</sub> (Deviance explained) |  | 0.359 (38.6%) |  |  |  | 0.169 (19.6%) |  |  |  | 0.118 (15.4%) |  |  |  |
| N (p-value, full/null comp) |  | 319 (< 0.001) |  |  |  | 314 (< 0.001) |  |  |  | 319 (0.022) |  |  |  |

**Table 3:** GAMM results of behavioral changes (suckling, riding, body contact with the mother, independent foraging) in the older offspring around sibling birth (± 2 years). Binomial GAMMs on proportions of time per day and individual. ID: Individual. S-birth = sibling birth, “:” = interaction term. * before/after = before/after sibling birth. Green: Classic interaction term derived from a separate model calculation (see methods section). Statistics for year (categorical control variable) not shown for clarity.

| Reference |  | Nursing |  |  |  | Riding |  |  |  | Independent Foraging |  |  |  | Body contact with mother |  |  |  |
| --- | --- | --- | --- | --- | --- | --- | --- | --- | --- | --- | --- | --- | --- | --- | --- | --- | --- |
| *Factor Variables:* | Category | Est. | SE | t | p | Est. | SE | t | p | Est. | SE | t | p | Est. | SE | t | p |
| (Intercept) |  | -7.34 | 0.84 | -8.73 |  | -2.20 | 0.78 | -2.80 |  | 0.93 | 0.40 | 2.31 |  | -2.67 | 0.52 | -5.15 |  |
| males | females | 1.18 | 1.01 | 1.16 | 0.25 | 1.56 | 0.73 | 2.21 | **0.034** | 0.03 | 0.55 | 0.05 | 0.96 | 0.03 | 0.75 | 0.04 | 0.97 |
| after YS-birth* | before* | 0.03 | 0.99 | 0.03 | 0.97 | -1.85 | 0.52 | -3.54 | **<0.001** | -0.10 | 0.28 | -0.35 | 0.73 | 0.38 | 0.11 | 3.35 | **<0.001** |
| Year |  | - | - | - | - | - | - | - | - | - | - | - | - | - | - | - | - |
| *Smooth term variables:* |  |  |  |  |  |  |  |  |  |  |  |  |  |  |  |  |  |
| T-S-birth: males |  | 5.16 | 5.99 | 26.14 | **0.001** | 1.00 | 1.00 | 0.81 | 0.37 | 5.96 | 0.68 | 11.56 | 0.24 | 1.00 | 1.00 | 5.05 | **0.025** |
| T-S-birth: females |  | 7.62 | 8.10 | 24.85 | **< 0.001** | 1.00 | 1.00 | 8.42 | **0.003** | 7.31 | 8.09 | 31.27 | **<0.001** | 1.00 | 1.00 | 7.14 | **0.008** |
| T-S-birth: males | females | 1.00 | 1.00 | 0.06 | 0.81 | 1.00 | 1.00 | 3.25 | 0.071 | 6.47 | 7.14 | 7.09 | 0.66 | 1.00 | 1.00 | 0.02 | **0.89** |
| Age: males |  | 1.80 | 1.94 | 0.89 | 0.58 | 1.00 | 1.00 | 33.45 | **<0.001** | 3.30 | 3.46 | 6.94 | 0.10 | - | - | - | - |
| Age: females |  | 1.00 | 1.00 | 6.33 | **0.012** | 1.00 | 1.00 | 10.59 | **0.001** | 1.00 | 1.00 | 2.00 | 0.16 | - | - | - | - |
| Age: males | females | 1.00 | 1.00 | 0.66 | 0.42 | 1.00 | 1.00 | 5.98 | **0.015** | 3.76 | 3.95 | 12.05 | **0.025** | - | - | - | - |
| Daytime |  | 1.13 | 1.25 | 2.36 | 0.13 | 3.32 | 3.74 | 4.99 | 0.19 | 2.20 | 2.69 | 24.87 | **<0.001** | 3.96 | 4.00 | 453.8 | **<0.001** |
| Seasonal effect |  | 2.34 | 3.00 | 20.63 | **< 0.001** | 0.79 | 3.00 | 1.79 | 0.064 | 2.79 | 3.00 | 88.63 | **<0.001** | 2.67 | 3.00 | 70.81 | **<0.001** |
| *Random effects:* |  |  |  |  |  |  |  |  |  |  |  |  |  |  |  |  |  |
| T-S-birth per ID (smooth) |  | 13.69 | 73.00 | 30.82 | **< 0.001** | 23.66 | 69.00 | 174.1 | **<0.001** | 52.07 | 76.00 | 1992.0 | **<0.001** | 33 | 73.0 | 264 | **<0.001** |
| Age per ID (smooth) |  | 9.18 | 64.00 | 45.18 | **< 0.001** | 1.69 | 58.00 | 4.76 | 0.006 | - | - | - | - | 5 | 65.0 | 31 | **<0.001** |
| Mother ID (intercept) |  | 0.00 | 10.00 | 0.00 | **< 0.001** | 0.00 | 10.00 | 0.00 | 0.019 | 4.14 | 12.00 | 7.61 | **<0.001** | 0 | 10.0 | 0 | **<0.001** |
| Date (intercept) |  | 0.00 | 1.00 | 0.00 | 0.19 | 0.00 | 1.00 | 0.00 | 0.36 | 0.00 | 1.00 | 0.00 | **<0.001** | 0 | 1.0 | 0 | 0.065 |
| R<sup>2</sup><sub>adj</sub> (Deviance explained) |  | **0.362 (59.3%)** |  |  |  | **0.830 (81.4%)** |  |  |  | **0.380 (44.4%)** |  |  |  | **0.317 (39.3%)** |  |  |  |
| N (p-value, full/null comp) |  | **545 (< 0.001)** |  |  |  | **301 (< 0.001)** |  |  |  | **471 (< 0.001)** |  |  |  | **545 (0.002)** |  |  |  |

### Physiological changes in response to transition to siblinghood (TTS) Urinary cortisol level changes in response to TTS

In the older offspring, we found at the time of a sibling birth a significant, sudden, non-continuous increase in urinary cortisol levels to about fivefold of the level prior to this event (S1A Fig.).

Further allowing for moderation of the TTS effect by the age of the older offspring (by adding respective interaction terms, see methods section) did not further improve the model, suggesting that the cortisol level changes were stable and not moderated by offspring age (Fig. 1B; Chi^2^(6) = 0.004, p = 1.00). Hence, the effect of TTS did not attenuate with increasing age of the older sibling (Fig. 1B).

**Fig. 1.**
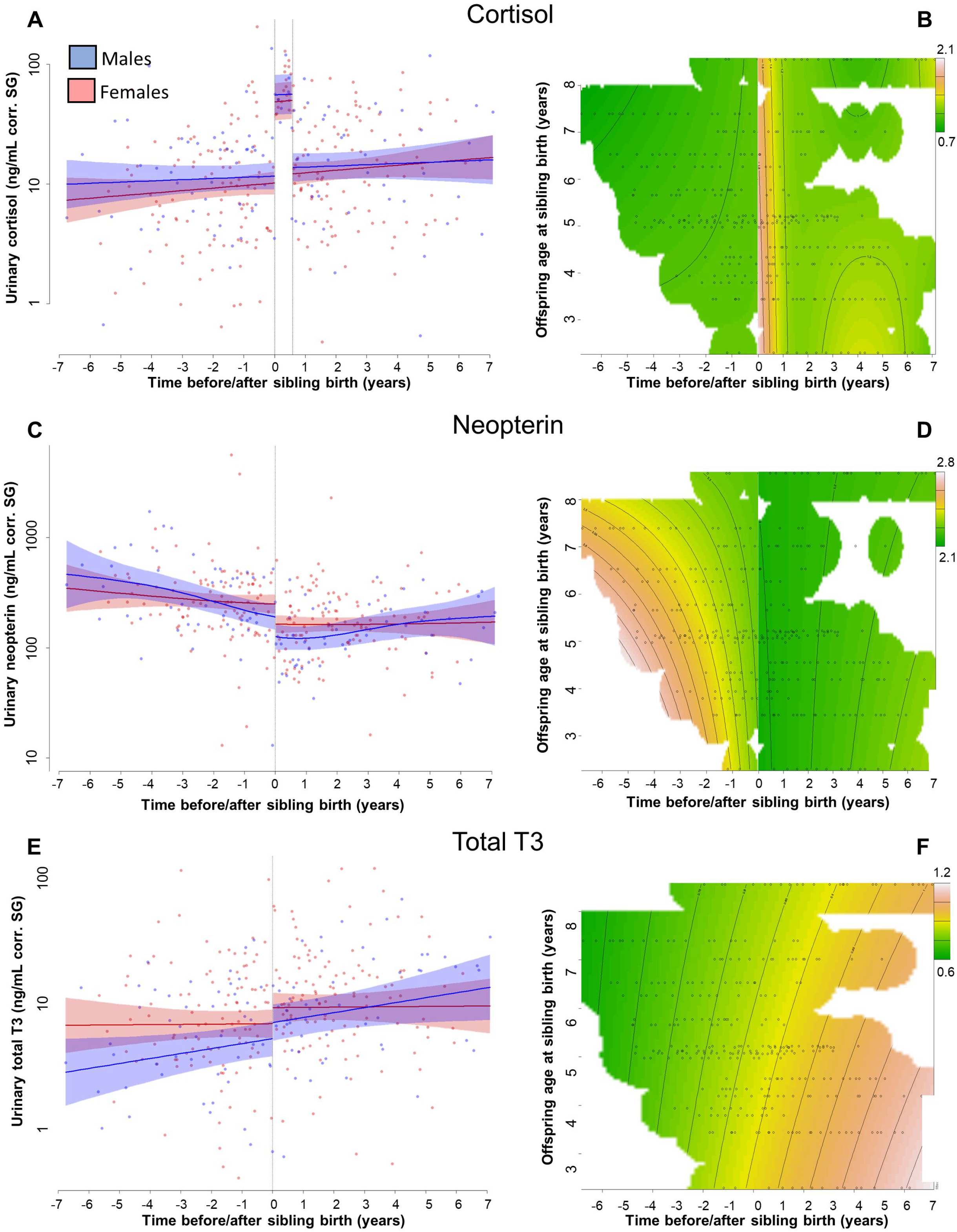
A-F: Physiological changes in the older offspring around sibling birth. Left: Main models from Table 1, right: Interaction plots further allowing for moderation of the TTS effect (left plot) by offspring age. Data points are physiological measures corrected for specific gravity (SG). All smooths uncontrolled for age to show cumulative pattern. **(A, B)** Urinary cortisol levels showed a significant, sudden rise to 10 ^0.67^ = 4.7-fold values at sibling birth (note the log scale) and remained at this level for seven months (right dotted line; no sex differences or age effects). **(C, D)** Urinary neopterin levels decreased by 1/3 at sibling birth (10^-0.19^ = 0.65; no sex differences or age effects). **(E,F)** Urinary total T3 levels increased around sibling birth.

Further post-hoc tests indicated that urinary cortisol remained at constant high level for about seven months after sibling birth and quickly decreased thereafter (Fig. 1A, B, Table 2). Data inspection revealed that cortisol measures in all samples collected within seven months following sibling birth were far above the upper 99.9% confidence interval of the values from before and after the 7-month post-birth period (Fig. S1A). To verify this pattern, we run another model allowing for an additional sudden change in cortisol levels not only right at sibling birth but also seven months later. This model significantly improved model fit (Fig. 1A; compared to the model with only one sudden change at sibling birth (Fig. S1B): Chi^2^(1) = 17.71, p < 0.001). Cortisol levels during the seven-month period after sibling birth were 10^0.67^ = 4.7 times higher than before sibling birth and 10^0.60^ = 4.0 times higher than after the seven-month period (Fig. 1A, Table 2). Cortisol levels after the seven-month period were not different from before sibling birth (Fig. 1A, Table 2). We found no significant effect of sex (Fig. 1A, Table 2).

### Urinary neopterin level changes in response to TTS

We found a significant sudden decrease to 65% (10^−0.19^ = 0.65) in urinary neopterin levels at the time of sibling birth (Fig. 1C; Table 2). Allowing moderation of this effect by the age of the older offspring did not improve the model (Fig. 1D; Chi^2^(6) = 0.78, p = 0.96). We found no sex effect (Fig. 1C, Table 2). With increase age urinary neopterin levels significantly changed in males but not females, but this sex difference was not significant (Table 2).

### Total T3 levels during TTS

Urinary total T3 levels increased with time around sibling birth (Fig.1E,F), but this pattern could not be clearly attributed to either age or sibling birth: the model including only time around sibling birth but not age was significantly better than the null model (p = 0.022, Table 2) whereas the model including both was not significantly different from the null model anymore (p = 0.142). There was neither a significant sex effect in urinary total T3 levels during TTS nor a significant sudden change in total T3 levels at sibling’s birth (Fig. 1E, Table 2). Adding interaction terms with age of the older offspring did not further improve the model (Fig. 1F; Chi^2^(6) = 1.83, p = 0.72).

### Behavioral changes in response to transition to siblinghood (TTS) Suckling in response to TTS

The proportion of time the older offspring was observed suckling showed a continuous decrease prior to sibling birth in both males and females, and reached zero about two months before parturition (Fig. 2A, Table 3). Consequently, there was no sudden change at sibling birth in offspring suckling (Fig. 2A, Table 3). There was no sex difference and suckling behavior declined significantly with age in females but not males (sex difference not significant, Fig. S2A, Table 3). Allowing for moderation by the age of the older offspring did not improve the model (Fig. 2B; Chi^2^(6) = −0.56, p > 0.1).

**Fig. 2:**
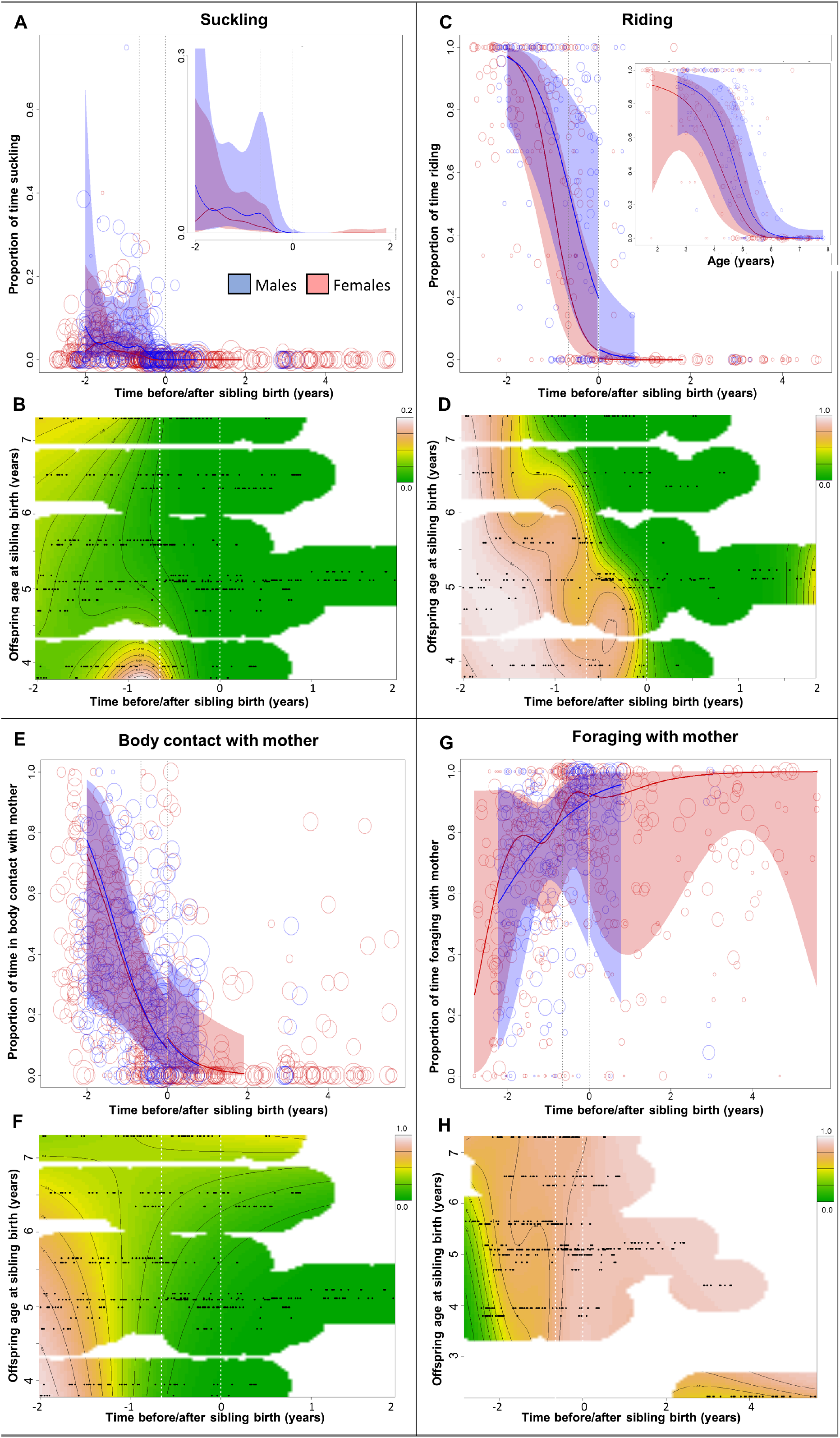
Behavioral changes in suckling, riding, body contact with the mother, and independent foraging of the older sibling in relation to sibling birth. Top and bottom plots, respectively: Main models from Table 2 (top) and interaction plots further allowing for moderation of the TTS effect by offspring age (bottom plot). Vertical dotted lines = time of putative conception and sibling birth. Data points are original proportion data per day and individual, with circle size representing the corresponding sample sizes (square-rooted; Ranges: Suckling and Body contact with mother 3 - 303, Riding 3-44, Foraging 1 - 182). All smooths uncontrolled for age to show cumulative pattern. **(A, B)** The proportion of time spent suckling decreased over time and ceased about two months before giving birth (**A**: Upper-right corner: zoom-in plot). **(C,D)** The proportion of time spent riding decreased towards and showed a sudden decline at sibling birth (**C**), but this sudden decline occurred only up to 5yrs of age (**D**). (**E,F**) The proportion of time spent in body contact with the mother declined largely before but also after sibling birth, with a slight sudden increase at sibling birth (**E**), but this effect attenuated with increasing offspring age (**F**). (**G,H**) Rates of independent foraging during maternal foraging bouts increased towards sibling birth but high independent feeding rates were already achieved long before maternal reconception.

### Riding on the mother in response to TTS

The proportion of time when the older offspring was riding on the mother during travel continuously decreased before sibling birth, then showed a significant sudden decline at sibling birth, and continued on very low levels afterwards (Fig. 2C, Table 3). Males spent significantly more time riding, and the continuous decline before sibling birth was only significant in females whereas the sudden decline at sibling birth seemed to be stronger in males (Fig. 2C, Table 3).

Allowing for moderation of the TTS effect by the age of the older offspring significantly improved the model (Chi^2^(6) = 10.25, p = 0.002), and visual inspection showed that the sudden decline in riding at sibling birth was only evident in young immatures whereas individuals older than five years of age were completely independent from maternal carrying long before sibling birth (Fig. 2D, Fig S2B). Hence, the effect of TTS on riding attenuated with increasing age of the older sibling, and disappeared in offspring older than five years.

### Spatial proximity to the mother in response to TTS

There was no effect of TTS on spatial proximity to the mother (within five meter), and none of the models was significantly different from the respective null models, indicating no consistent effect beyond random and temporal variation in our data set.

### Body contact with the mother in response to TTS

The proportion of time the older offspring spent in body contact with the mother decreased before and around sibling birth, reaching low levels already during gestation (Fig. 2E, Table 3). However, this pattern could not be clearly attributed to either age or time around sibling birth, and the model including both variables was not significantly different from the respective null model (p > 0.1; but see also Fig. 2F) whereas the model without age was significantly different (p = 0.002, Table 3). There was a small but significant sudden change at sibling birth, but in contrast to the continuous decline expected for weaning, this was a sudden increase of body contact with the mother (Fig. 2E, Table 3). Allowing moderation of the TTS effect by the age of the older offspring significantly improved the model (Chi^2^(6) = 75.33, p < 0.001), and visual inspection showed that body contact decreased with age of the older sibling, with the consequence that also the decline in body contact before sibling birth attenuates with age (Fig. 2F).

### Independent foraging in response to TTS

The proportion of time that older offspring spent foraging independently while the mother was foraging, significantly increased towards sibling birth, and this effect was moderated by the older offspring’s age (Fig. 2G, Table 3; allowing moderation of the effect by the age of the older offspring: Chi^2^(6) = 15.58, p < 0.001, Fig. 2H). However, these changes occurred mainly before conception of the mother, and all subjects were almost entirely independent in terms of foraging at the time of sibling birth, irrespective of their age, and there was no sudden change around sibling birth (Fig. 2H, Table 3).

## Discussion

Our data from wild bonobos demonstrate that sibling birth induces a physiological stress response in the older offspring that is independent of the older offspring’s age. Upon birth of a sibling, urinary cortisol levels in the older offspring increased fivefold and remained at this level for about seven months. Parallel to this change in urinary cortisol levels, urinary neopterin levels declined at sibling birth, indicating reduced cell-mediated immunity. These physiological changes around sibling birth did not attenuate with offspring age and were independent of our measures of nutritional and social weaning. At sibling birth, weaning-related changes were either already completed (independent foraging and suckling), did not show a sudden change (urinary total T3, suckling, independent foraging, and time in spatial proximity to mother), or were only a significant factor at younger ages but ceased with increasing offspring age (riding and body contact time with the mother).

The five-fold increase in cortisol that we observed in our study is an unusually strong physiological response. For comparison, captive bonobos exposed to an experimental stress test exhibited a two-fold increase in cortisol levels (Verspeek et al., 2021). A similar cortisol response occurs in group members in response to infant birth, but levels returned to normal within one day (Behringer et al., 2009). In wild chimpanzees, urinary cortisol levels were found to increase by a factor of 1.5 when subjects encounter a neighboring group, an event that exposes all group members to potentially lethal aggression (Samuni et al., 2019). Cortisol level changes exceeding what we found in our study is a ten-fold increase in wild chimpanzees suffering from a respiratory disease that killed many individuals (Behringer et al., 2020), and probably also disrupted many social ties among those who survived. The intensity of a stress response is generally determined by the severity, controllability, and predictability of the stressor (Seiler et al., 2020), all of which probably apply to TTS as a stressor and contribute to the comparably high cortisol response that we observed in our study.

In addition to the age-independent, sudden, and substantial physiological stress response that we observed, we found that cortisol levels remained elevated for seven months after sibling birth, even though nutritional and social weaning was largely completed by this time. In chimpanzees, anecdotal reports indicate that it may take one year until the older offspring adapts behaviorally to the conditions following sibling birth (Clark, 1977). While the physiological effects of sibling birth in human children are still unknown, behavioral data suggest that it may take up to eight months until the older sibling adapts to the novel situation (Oh et al., 2017; Stewart et al., 1987). This indicates that humans and apes are similar in their response to the challenge deriving from the arrival of a sibling.

Cortisol levels increase in response to psychological and social stressors, like predation risk or social instability, as well as an energetic and physiological stressors (McEwen and Karatsoreos, 2020). Our study indicates that the sudden and persistent increase in cortisol levels in the older sibling was not related to an energetic stressor. Neither urinary total T3 levels, nor suckling frequency, nor time spent foraging independently from the mother showed abrupt changes at the onset of TTS. If the physiological stress response at sibling birth was caused by energetic challenges, the intensity of the cortisol level change would decline with the age of the older offspring due to decreasing dependency of the mother. The age of the older offspring at sibling birth in our study ranged from 2.3 to 8.6 years, and did not affect cortisol level changes. Preliminary analyses of stable isotopes in our study group suggest that nutritional weaning is completed at around 4.5 years of age (Oelze et al., 2020), and females disperse at the age of eight years which indicates independence from maternal support at this age (Furuichi, 1989; Hashimoto, 1997; Kuroda, 1989). The behavioral profiles (body contact, suckling, and independent foraging) of the immatures involved in our study showed a gradual change of dependency with age instead of a sudden shift at the time of sibling birth. These findings emphasize that the sudden increase in cortisol levels is independent from other weaning effects, resembling findings from human children showing behavioral responses to sibling birth independent of their actual age (Dunn and Kendrick, 1980; Stewart et al., 1987). Hence within the scope of our behavioral metrics, cortisol patterns did not match changes in single or cumulative behavioral changes around sibling birth or throughout siblinghood.

The stress response of older offspring at sibling birth is may be associated with changes in their social environment, such as reduced social support from mothers and the concomitant exposure to novel social challenges that characterize maturation (Dunn and Kendrick, 1980; Kendrick and Dunn, 1980). For example, in wild bonobos, juveniles of both sexes are exposed to severe physical aggression from adult males, and aggression rates increased with age of the target (average: 5.9 years of age; range 3-7.9 years of age). In contrast, adult males bonobos never directed aggression against infants (Hohmann et al., 2019). The age when young bonobos start becoming targets of adult male aggression matches with the age at TTS.

Although body contact between the older offspring and the mother decreased with age, it also suddenly increased for a short period after sibling birth. Similarly, juvenile marmosets increased proximity to parents after sibling birth (Achenbach and Snowdon, 1998), infant rhesus macaques increased their effort to maintain contact with their mothers (Mandalaywala et al., 2014), and children increase clinging behavior during TTS (Volling et al., 2017). In our study, we did not find consistent effects of TTS on proximity (within five meters). If such changes in proximity and body contact reflect reduced maternal attention, older offspring may aim to regain more attention from their mothers / care givers (Baydar et al., 1997). Reduced maternal attention could contribute to the increase in cortisol levels that we found, but it is still unclear why this change persists for several months. Moreover, the most consistent effect of TTS on offspring behavior in humans was a decrease in affection and responsiveness to the mother (Volling, 2012), which seems to counteract this interpretation. Alternatively, an increase in body contact could also be interpreted as curiosity and interest in the baby, as young female primates show a high interest in new babies independent of age and kinship (Maestripieri and Pelka, 2002).

The sudden increase in cortisol and the abrupt decline in neopterin levels in our study emphasizes the homeostatic challenges affecting older offspring during TTS. It is possible that the increase in cortisol levels negatively affected cell-mediated immunity. In other mammals, stress responses to weaning had a negative effect on immunity (Kick et al., 2012; Kim et al., 2011), and stressful events are associated with changes in immune function in humans (Herbert and Cohen, 1993). While short-term increases in cortisol levels enhance immune functions in humans, long-lasting elevations of cortisol levels—such as those that we found in our study — dysregulate immune responses (Dhabhar, 2014). In our study, urinary cortisol and probably also neopterin levels recovered several months after sibling birth, indicating that individuals can cope with TTS to some degree, for example by becoming habituated to the new conditions or by recruiting social support from other group members.

Persistent early-life stress can affect an individual’s ontogeny with long-lasting consequences for its fitness, affecting its growth trajectory, metabolism, social behavior, immunity, stress reactivity, reproduction, and life history strategies (Berghänel et al., 2017; Maestripieri, 2018; Seiler et al., 2020). In view of our results, such effects may contribute to the observed negative effects of sibling birth on the fitness of the older offspring in non-human primates (Emery Thompson et al., 2016; Tung et al., 2016; Zipple et al., 2019). However, strong negative effects caused by a highly predictable and normative stressor that invariably affects most individuals should be under negative selection. It is tempting to speculate that this pronounced early-life stressor is “tolerable stress” (McEwen and Karatsoreos, 2020) that may prime subjects to develop stress resistance later in life. Furthermore, TTS can relate to accelerated acquisition of motor, social and cognitive skills (Azmitia and Hesser, 1993; Maestripieri, 2018; Song et al., 2016). Siblings are not only stressful rivals but also important social partners, and the presence of an older sibling can buffer behavioral and physiological changes in response to stressful events like TTS (Hrdy, 2011). Having an older sibling may enhance the development and survival of the younger sibling which contribute to the inclusive fitness of both the older sibling and the mother (Salmon and Hehman, 2015; Stanton et al., 2017).

Our study on wild bonobos, is to our knowledge, the first to investigate the physiological stress response during TTS. In addition, as other studies on nonhuman primates, it sheds light on the evolutionary origins of patterns of TTS. In many human cultures, inter-birth intervals are shorter and children are weaned at a younger age than in wild apes (Humphrey, 2010; Robson et al., 2006). However, parental effort varies tremendously across human cultures and is often supplemented by intense allomaternal care (Hrdy and Burkart, 2020). Thus, it is possible that children do not typically experience such extreme and long-lasting cortisol fluctuations. In some families of western societies and traditional societies, allomothering provides nutritional, physical, and mental support to older children (Baydar et al., 1997; Kramer and Veile, 2018), possibly buffering physiological stress responses. However, when such social buffering systems are absent or weakly developed as perhaps in some modern and particularly western societies, older children may experience the birth of a sibling as a particularly stressful time. Since studies in humans are generally biased towards middle-class families in western industrialized countries, the current evidence does not necessarily allow to make generalizations about humans (Fouts and Bader, 2016; Volling, 2012). Our study on wild bonobos contributes to this discussion by showing that TTS is stressful also under natural conditions, highlighting its evolutionary history.

Altogether, our results on wild bonobos support the long-standing but untested and recently questioned assumption that the birth of a sibling is a highly stressful event for the older offspring (Volling, 2012; Volling et al., 2017). It highlights the ubiquity of this pattern across individuals and age classes, and indicates its evolutionary roots. However, behavioral responses to TTS in human children are highly variable and individual- and age-dependent, and range from aggression, emotional blackmailing, and psychological disturbances to positive attitudes towards the new constellation and co-parenting (Volling, 2012; Volling et al., 2017). This raises questions regarding the coping strategies and how they are a) influenced by the socioecological conditions including actual parent-offspring and other caretaker relationships, b) effective in modulating and buffering the shown physiological stress response, and c) unique to humans or do also occur in non-human primates and other animals (Hrdy, 2011; Lonsdorf et al., 2018).

## Methods

### Study site and species

Data were collected from wild bonobos (*Pan pansicus*) of the Bompusa West and East communities, at LuiKotale, Democratic Republic of the Congo. This bonobo population was never provisioned and lives in an intact natural forest habitat. All subjects were habituated to human presence before the start of the study, were genotyped, and were individually known. We considered every offspring only for the next sibling birth, therefore, all older offspring in our study experienced the birth event for the first time. At the birth event, the older siblings were between 2.3 and 8.6 years old. The behavior sampling included 397.17 hours of focal data on eleven immature females (Mean = 36.11, SD = 14.70) and 253.95 hours on six immature males (Mean = 42.33, SD = 27.62). Physiological measurements were performed in 319 (220 female, 99 male) urine samples of 20 females and six males.

### Behavioral data collection and analysis

Behavioral data were collected systematically between July 2015 and July 2018 via focal animal sampling (Altmann, 1974) whereby an infant was observed for one hour and its instantaneous behavior recorded at one-minute intervals. Behaviors included suckling, defined as the infant applying its mouth to the nipple of the mother in a suckling manner, and riding, defined as the infant being transported as it clings ventrally or dorsally to its mother. We recorded when the offspring was in body contact or within 5-meter proximity to the mother. For riding, we only considered data where the mother was travelling for at least three consecutive minutes to exclude situations where the mother was likely travelling for short distances only and riding on the mother would not have been important for the offspring. For independent foraging of the offspring, we only considered scans where also the mother was foraging to cover typical foraging situations and reduce the influence of potential sampling bias, with foraging encompassing handling and ingesting food.

### Urine sample collection and analyses

Urine samples for physiological analyses were collected between July 2008 and August 2018. Urine samples were collected opportunistically throughout the day between 5 am and 6 pm. Samples were collected from the vegetation or directly into leaves. Samples contaminated with feces were excluded. All other samples were protected from direct sunlight to avoid degradation, carried to camp, and frozen in liquid nitrogen upon arrival on the same day. Samples were shipped frozen to the Max Planck Institute for Evolutionary Anthropology in Leipzig, Germany, for cortisol and total triiodothyronine analysis, and later to the German Primate Center, Göttingen, Germany for neopterin measurement.

Our urine data set consists of 16.0 +− 5.6 samples per individual (mean +-SD), with on average 7.5 samples before and 8.4 samples after sibling birth. Urine samples were temporarily normally distributed around the day of sibling birth. Urine samples were collected from all individuals also during the first year after sibling birth, though one male and two females did not contribute samples during the first seven months after sibling birth, and therefore, contributed only to the estimates of the urinary cortisol levels before and after the elevated cortisol period (results section).

Frozen samples were first thawed at room temperature, shaken for 10 seconds (VX-2500 Multi-tube Vortexer) and centrifuged for 5 minutes at 2.000 *g* (Multifuge Heraeus), after which specific gravity (SG) was measured using a refractometer. All results were corrected for SG, to adjust the concentration of the physiological marker for urine concentration of the specimen, which depends on an individual’s hydration status and time since last urination (Miller et al., 2004). Aliquots of samples were prepared at this time for later neopterin and total T3 analyses.

#### Urinary cortisol analyses

We extracted and measured urinary cortisol in 319 (220 female, 99 male) urine samples of 20 females and 6 males with liquid chromatography–tandem mass spectrometry (LC-MS/MS) and MassLynx (version 4.1; QuanLynx-Software). Cortisol extraction from urine samples was performed following the protocol described in Hauser et al. (2008) for liquid chromatography–tandem mass spectrometry analyses. Each urine sample was mixed with an internal standard (prednisolone, methyltestosterone, d3-testosterone, d4-estrone and d9-progesterone). Internal standard was used as quality control to assess sample recovery and to quantify urinary cortisol levels using prednisolone. We performed hydrolysis using β-glucuronidase from *Escherichia coli* (activity: 200 U / 40 μl). Extracts were purified by solid phase extractions (Chromabond HR-X SPE cartridges: 1 mL, 30 mg). Followed by a solvolysis with 250 ml ethyl acetate and 200 mg sulphuric acid. The extraction of cortisol was carried out with methyl *tert*-butyl ether. Finally, we reconstituted evaporated extracts in 30% acetonitrile.

For urinary cortisol measurement we used a liquid chromatography-tandem mass spectrometry (LC-MS/MS) with a Waters Acquity UPLC separation module equipped with a binary solvent manager, sample manager, and a column oven (Waters, Milford, MA, USA). Waters Acquity BEH C18 column (2.1 x 100 mm, 1.7 μm particle diameter) were used for chromatographic separation. Eluent A was water with 0.1% formic acid and Eluent B was acetonitrile. We injected 10 μl. We performed mass spectrometry analyses on a Waters XEVO TQ-S tandem quadrupole mass spectrometer (Waters, Milford, MA, USA) with an electro spray interface (ESI) in positive mode. The quantitative analysis of cortisol levels was realized in the range of 0.01–100 pg/μl. For cortisol examination we used MassLynx (Version 4.1; QuanLynx-Software). Final urinary cortisol results are represented in ng/ml corrected for SG. We accepted measurements of a batch if quality control measurements deviated less than 13 % from the true cortisol concentration. 17 samples in which internal standard recovery deviated by more than 60% of the internal standard were re-measured via reinjection. In two samples, measurements were above the limit of the calibration curve, we reinjected extracts at a 1:10 dilution.

#### Urinary neopterin analyses

We measured urinary neopterin in 314 (215 female, 99 male) aliquots of 20 females and 6 males with a commercial neopterin ELISA for humans, previously validated to determine neopterin in bonobo urine (Behringer et al., 2017). Prior to neopterin measurement, urine samples were diluted (1:10–1:200 depending on SG) with the assay puffer provided by the supplier. We added to each well on the plate 20 μl of the diluted urine, 100 μl of the provided enzyme conjugate, and 50 μl of the neopterin antiserum. The plate was covered and incubated on an orbital shaker at 500 rpm in the dark for 90 minutes. The plate was then washed four times with 300 μl washing buffer, and 150 μl of tetramethylbenzidine substrate (TMB) solution was added. The plate was incubated again for 10 minutes and the reaction was stopped by adding150 μl of the provided stop solution. Optical density was measured photometrically at 450 nm.

All samples were measured in duplicates according to the supplier’s instructions. Inter-assay variation for high- and low-value quality controls was 4.2 and 1.7 % (N = 17 assays), respectively. Intra-assay variation was 8.9 %. Final neopterin concentrations are expressed in ng/ml corrected for SG.

#### Urinary total T3 analyses

We measured total T3 in 319 (220 female, 99 male) urine aliquots of 20 females and 6 males with a commercial, competitive total triiodthyronine (T3) ELISA (Ref. RE55251, IBL International GmbH, Hamburg, Germany). Samples were measured with a 1:2, 1:5 or without dilution depending on SG. 50 μl of the diluted sample with 50 μl of the provided assay reagent was pipetted into a well. We shanked the plate for 10 seconds and incubated the plate afterwards for 30 minutes at room temperature. We then added 50 μl of the provided Triiodothyronine-enzyme conjugate to each well, shacked the plate again for 10 seconds and incubated it again at room temperature for 30 minutes. We then washed the plate five times with 300 μl of the washing puffer and added 100 μl of TMB substrate. After 10 minutes of incubation, we stopped the reaction with 100 μl of the provided stop solution and read the plate at 450 nm with a microplate reader.

All samples were also measured in duplicates. Inter-assay variation for high-and low-value quality controls was 6.3 and 5.6 % (N = 25 assays), respectively. Intra-assay variation was 7.2%. Final total T3 concentrations are expressed in ng/ml corrected for SG.

### Statistical analysis

All statistical analyses were performed with R 4.0.5 (R Development Core Team, 2020). We applied Generalized Additive Mixed Models (GAMM) which allow for the detection and analysis of complex non-linear relationships (termed “smooths”) that are typical for developmental trajectories. We used function gam for all models (package mgcv (Wood, 2017)), with smooth estimation based on penalized cubic regression splines. We checked for model assumptions and appropriate model settings using functions gam.check (package mgcv), and all models were inspected for and showed negligible auto-correlation (function acf_resid, package itsadug (van Rij et al., 2020)) and overdispersion (functions testDispersion and testZeroInflation, package DHARMa, (Hartig, 2021)). Model comparisons were conducted using the function compareML (package itsadug). GAMM smooths were plotted using package itsadug (van Rij et al., 2020) with removed random effects. As typical for GAMMs, interaction terms with factor variables were calculated in two ways, first analysing whether significant changes occur within each level of the grouping factor, and second whether the smooths of the different levels differ significantly from each other (the classic interaction term statistic) (Wieling, 2018; Wood, 2017).

Urinary physiological data (urinary cortisol, total T3, and neopterin) were normally distributed after log-transformation, and Gaussian GAMMs were applied. The GAMMs on mother-offspring relationship (suckling, riding, independent foraging, body contact, and proximity) were based on single minute-by-minute focal scan records which were summed to time proportion values per day and individual, hence we applied GAMMs with a binomial logit-link error structure on proportion data and the underlying number of scans per proportion value as weight-argument. The main predictor variable of all analyses was the temporal change of the respective response variable around sibling birth, allowing for potential sex differences (Behringer et al., 2014; Leigh and Shea, 1996).

Time around sibling birth was added in two ways into the model, first as a continuous smooth term across time, and second as a factor variable coding for the time before and after sibling birth, thereby allowing for a sudden, non-continuous and unconnected change right at sibling birth. Implemented this way together with the smooth term, this factor term directly estimates the sudden change at sibling birth and its error and statistical significance. Additionally, all models included potential mediating effects of age to control whether apparent TTS effects were in fact mere general age effects irrespective of TTS. Age and time around sibling birth were naturally 100% correlated within individuals and highly correlated within the entire datasets (range r = 0.659 - 0.855).

In a further step, we expanded these two terms of time around sibling birth to interaction terms incorporating offspring age at sibling birth, to investigate a potential moderation effect of offspring age on the intensity and pattern of potential TTS-effects. For this purpose, we run model comparisons of these interaction models with the respective models without interaction terms to estimate whether the implementation of a moderation effect of offspring age further contributed to model fit.

All statistical GAMMs were controlled for repeated measurements per individual via a) two random smooth effects (factor-smooth-interactions, for details see (Wood, 2017)), one for individual changes over time relative to sibling birth and the other for individual changes with age (for those models that included age as predictor variable), and b) a random intercept per mother since some mothers contributed multiple offspring. All GAMMs were controlled for year (as random intercept for hormonal data but as control variable for the three years of behavioral data), seasonal effects via a cyclic smooth term over the year, and for daytime effects via a smooth term over daytime. The binomial models on behavioral time proportion data included an additional random intercept of date to control for multiple measurements per day. Due to the structure of the interaction models combining age and time around sibling birth into one interaction term, we did not additionally control for general mediating age effect, and merged the random effects on age and mother ID to one random smooth term of age at sibling birth per mother ID.

## Ethics

All samples were collected non-invasively and with permission of the Institut Congolais pour la Conservation de la Nature (ICCN).

## Data accessibility

Source data for statistics and figures in the paper is permanently stored at GRO Behringer, 2021, “Replication Data for: Transition to siblinghood”, https://doi.org/10.25625/O1OD2I.

## Author contributions

Conceptualization: V.B., A.B., S.M.L. and G.H.; Methodology: V.B., A.B. and S.M.L; Software and Formal Analysis: A.B.; Validation: V.B.; Investigation: V.B., S.M.L., B.F. and G.H.; Resources: V.B, A.B., B.F. and G.H; Data curation: V.B, A.B., S.M.L., B.F. and G.H.; Writing – Original Draft: V.B., A.B. and G.H.; Writing –Review & Editing: A.B., V.B., S.M.L., B.F. and G.H.; Funding Acquisition: V.B., S.M.L., B.F. and G.H.; Supervision: B.F. and G.H.

## Acknowledgments

We thank the Institut Congolais pour la Conservation de la Nature (ICCN), and the people of Lompole for granting permission to conduct fieldwork on bonobos in their forest in the buffer zone of Salonga National Park. We are extremely grateful to all assistants of the LuiKotale Bonobo Project.

## Supplement figures legends

**Fig. S1.:**
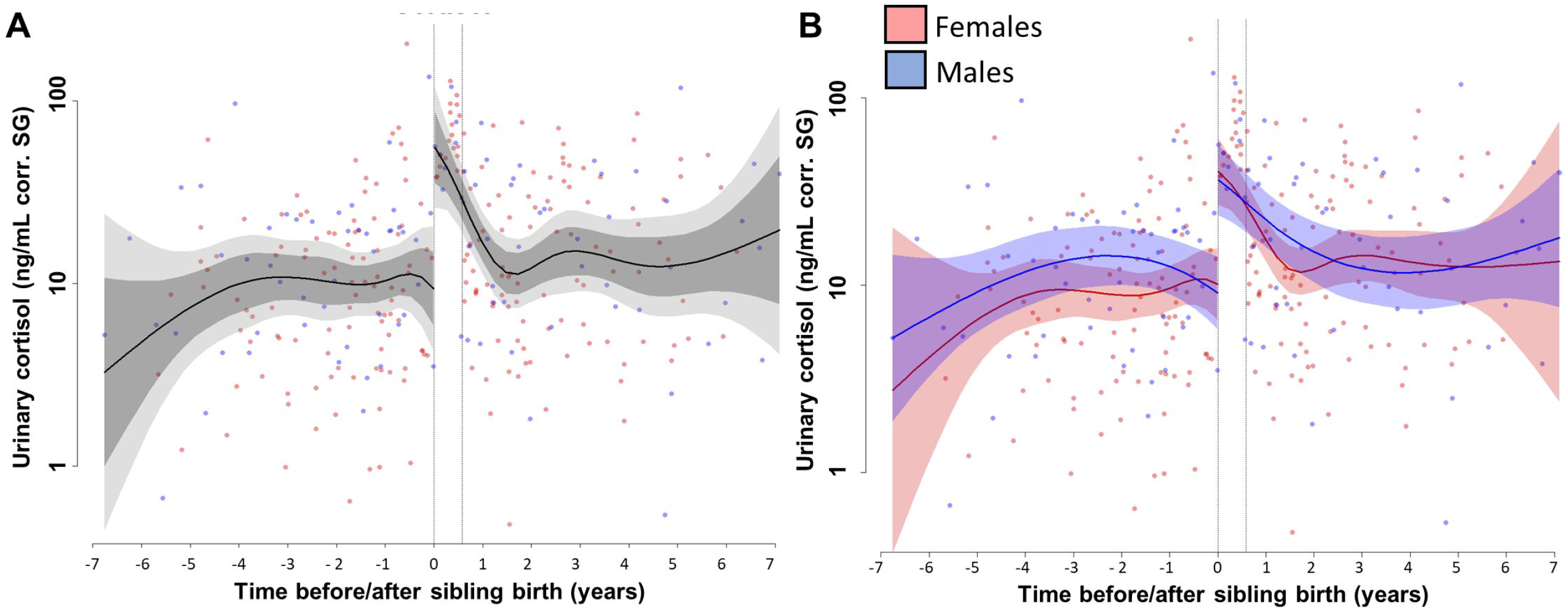
(A) A separate, independent smooth for urinary cortisol levels before and after sibling birth, allowing for an abrupt change (plot including 99.9% confidence intervals, light grey). (B) Same as (A) but with sex-specific trajectories. Data points are original data corrected for specific gravity. All smooths controlled for age. SG: specific gravity.

**Fig. S2.:**
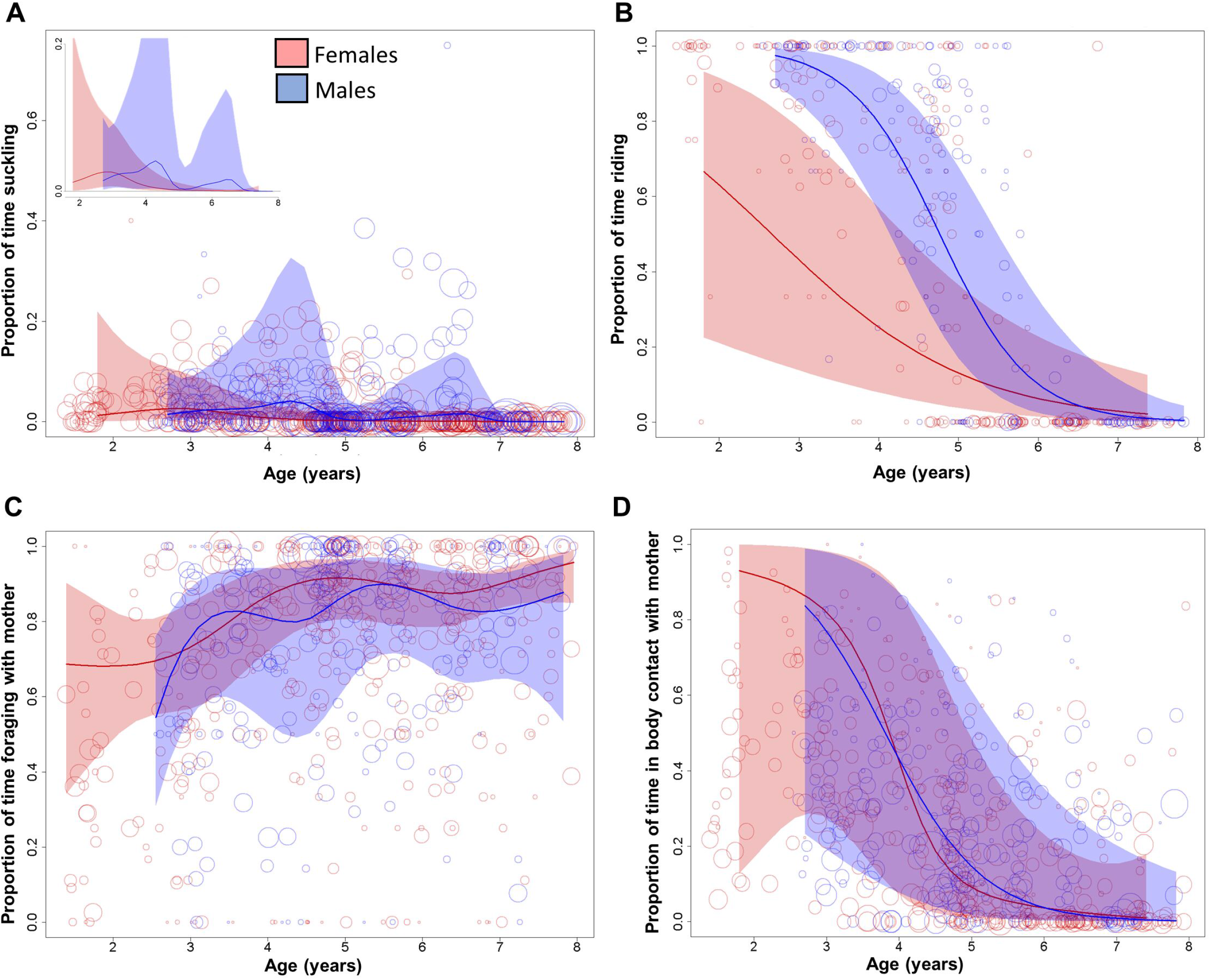
Behavioral changes in A) suckling, B) riding, C) independent foraging of the older sibling, and D) body contact with the mother in relation to sibling birth. Vertical dotted lines = time of putative conception and sibling birth. Data points are original proportion data per interaction bout, with circle size representing the corresponding sample size (square-rooted; Ranges: Suckling and Body contact with mother 3 - 303, Riding 3-44, Foraging 1 - 182). All smooths uncontrolled for age to show cumulative pattern.

